## Supplementary Figures for "A machine vision guided robot for fully automated embryonic microinjection"

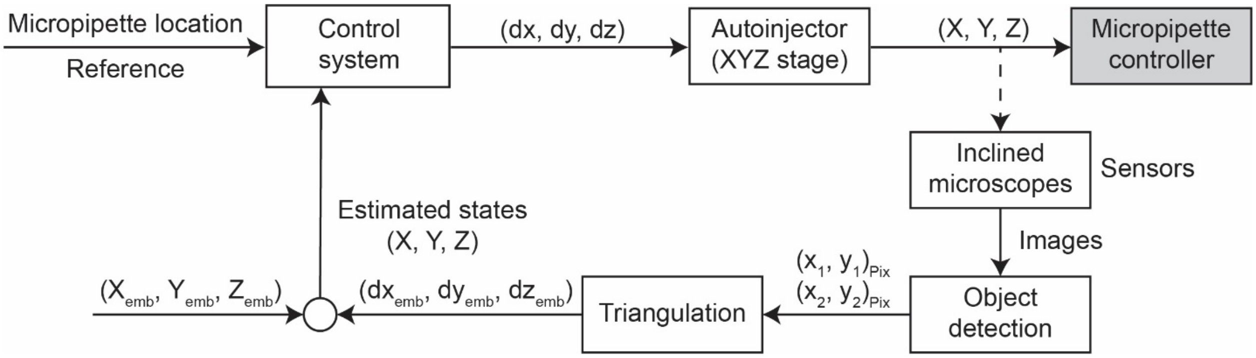


**Supplementary Figure 1: Procedure for microinjection location estimation:** Overall microinjection point estimation procedure for microinjection of Drosophila and zebrafish embryos.

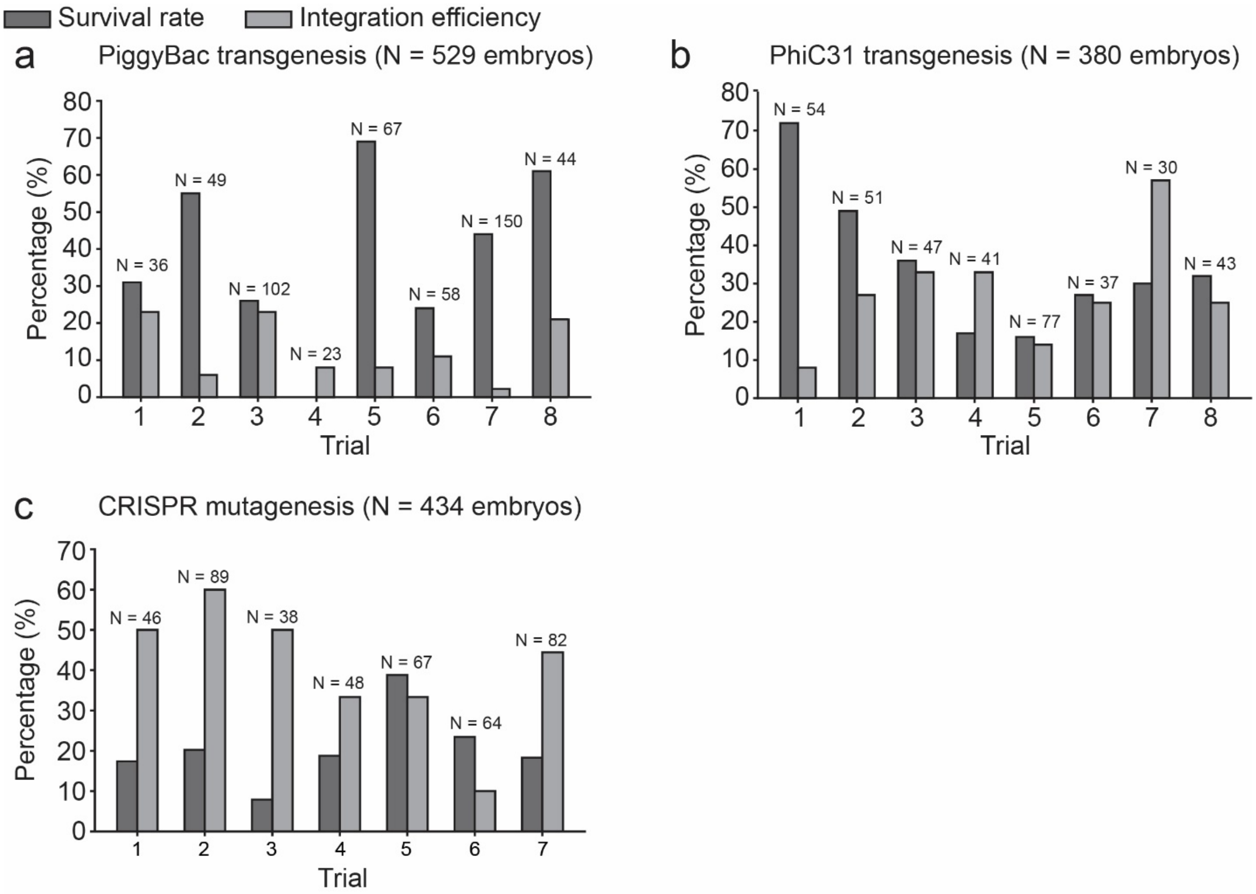


**Supplementary Figure 2: Trial-by trial breakdown of post-injection survival rate in germline transgenesis experiments conducted in Drosophila:** (a) PiggyBac transgenesis, (b) PhiC31 transgenesis, and (c) CRISPR mutagenesis. Number of embryos injected (N) are shown on the plots.

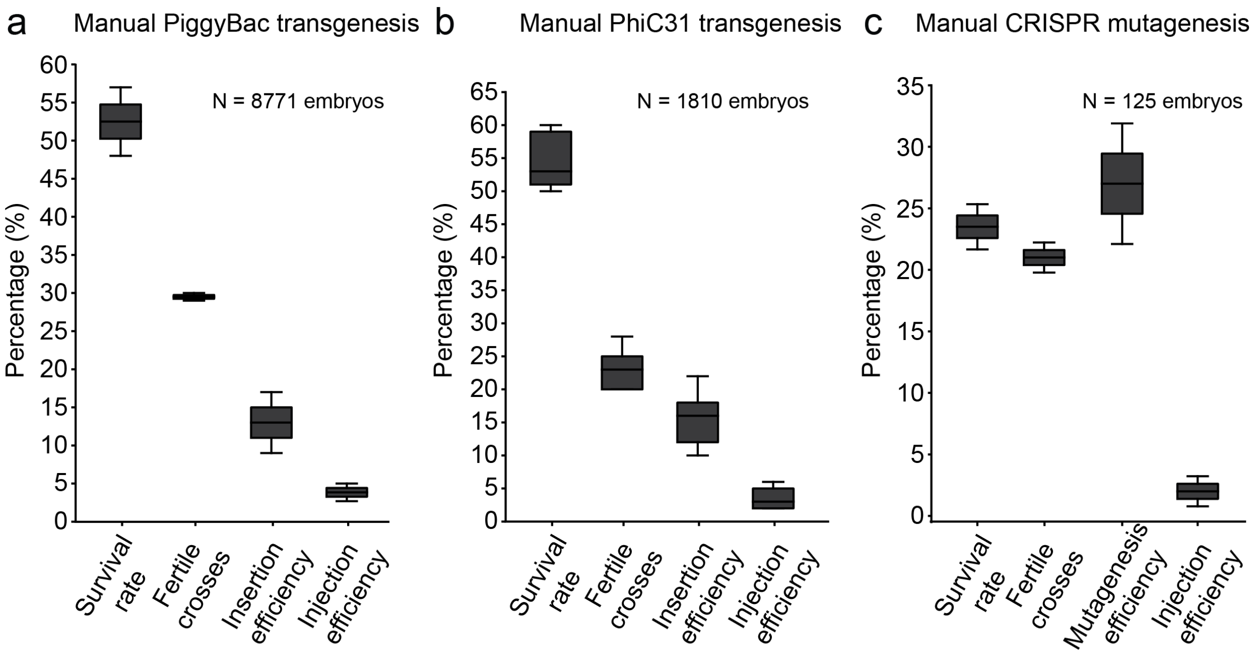


**Supplementary Figure 3: Post-injection survival rates, % fertile crosses, insertion efficiencies and injection efficiencies obtained with manual microinjections in Drosophila** (a) PiggyBac transgenesis, (b) PhiC31 transgenesis, and (c) CRISPR mutagenesis.

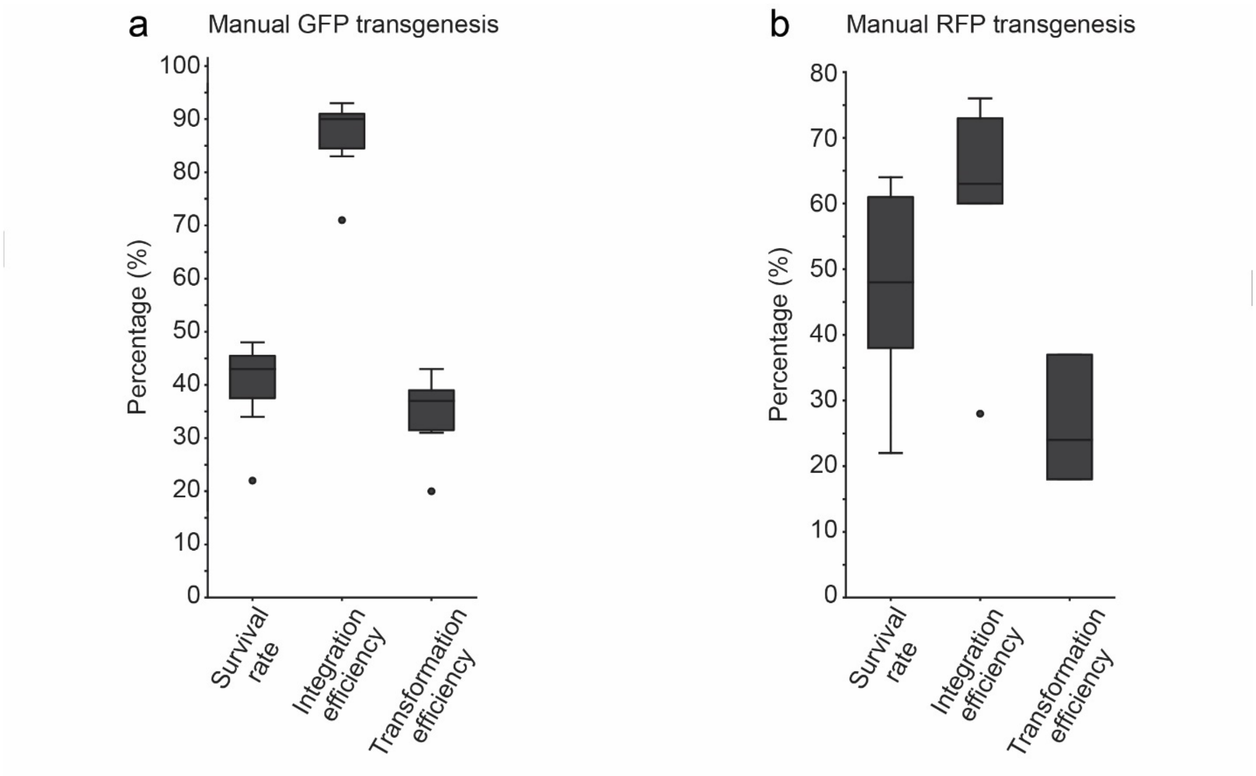


**Supplementary Figure 4: Post-injection survival rates, integration efficiencies and transformation efficiencies obtained with manual microinjections in zebrafish:** (a) GFP transgenesis and (b) RFP transgenesis.

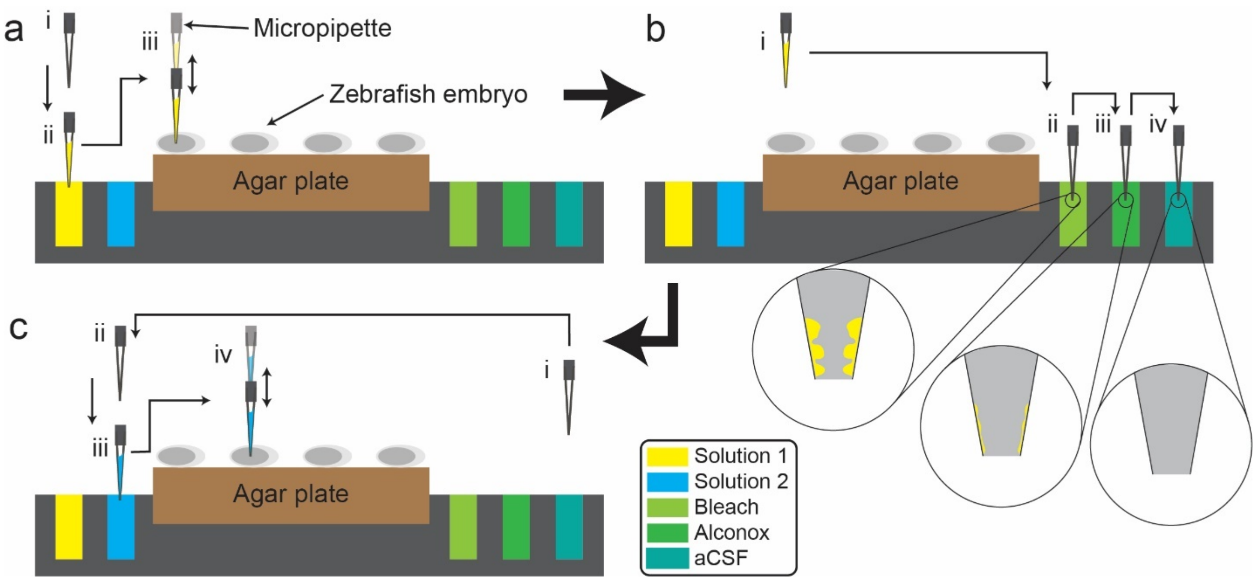


**Supplementary Figure 5: Multi-solution system:** Multi-solution system used for zebrafish embryo microinjections. (a) microinjection of an embryo with solution 1: i) Micropipette before micropipette front filling ii) front filling of first solution iii) microinjection of first solution (b) cleaning of micropipette: i) micropipette after microinjection of first solution ii) cleaning of micropipette via bleach, iii) Alconox, and iv) aCSF, and (c) microinjection of an embryo with solution 2: i) micropipette after cleaning ii) micropipette before micropipette front filling iii) front filling of second solution iv) microinjection of second solution.

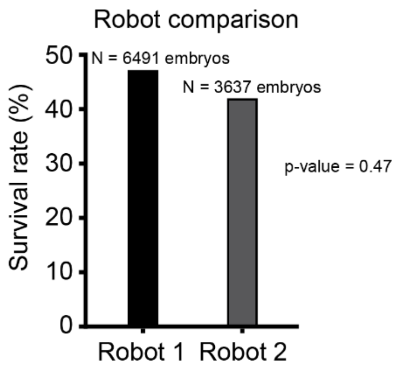


**Supplementary Figure 6: Robot comparison:** Larvae survival rate comparison of two independent microinjection robots used for Drosophila embryo microinjections. A two-sample t-test was used for statistical testing.

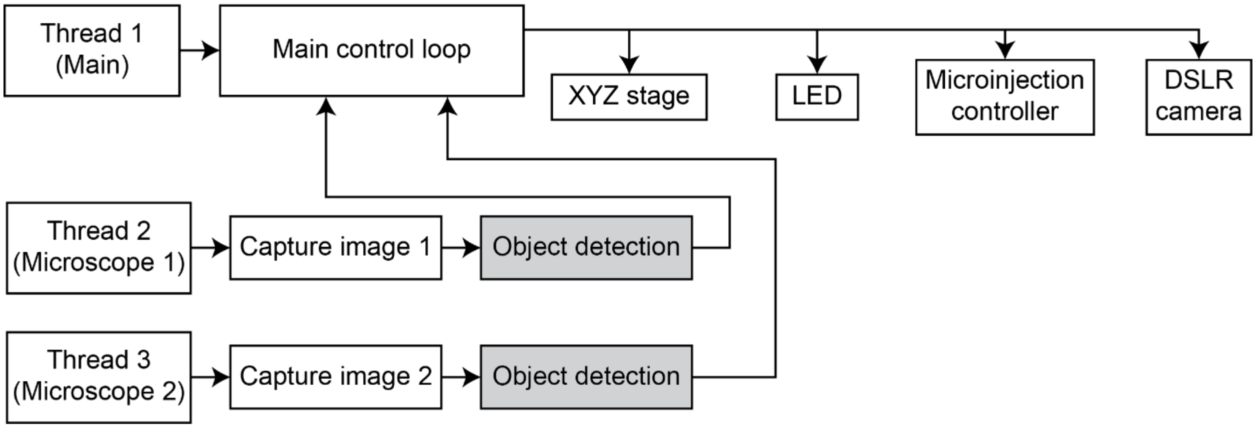


**Supplementary Figure 7: Software architecture for robot:** Overall software architecture used for automated microinjection of zebrafish and Drosophila embryos.

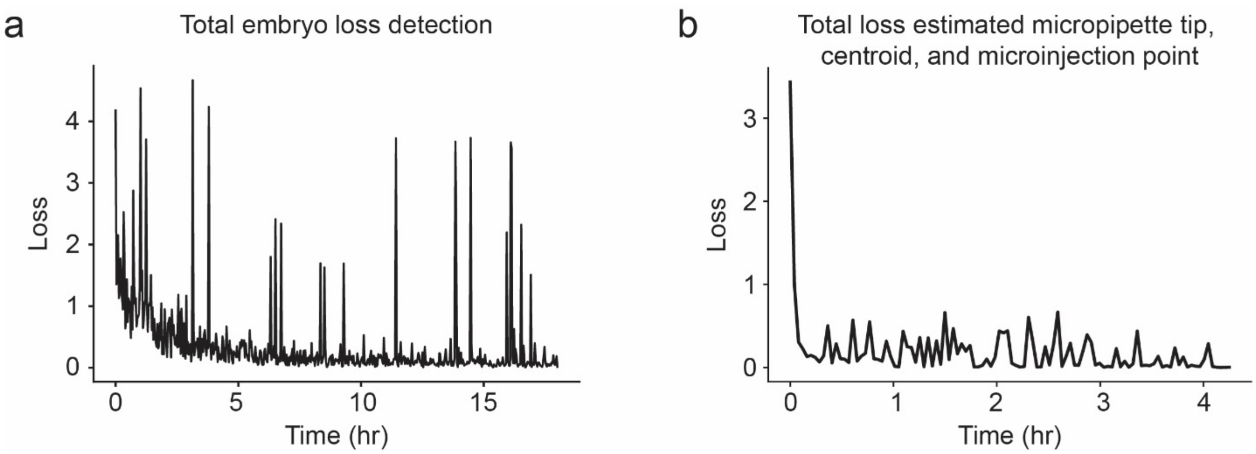


**Supplementary Figure 8: Machine learning loss rate for Drosophila dataset training:** (a) ML loss rate for the model used to detect embryos at macroscale via the DSLR image. (b) ML loss rate for the model used to detect the micropipette tip, centroid of the embryo, and microinjection point of the embryo.

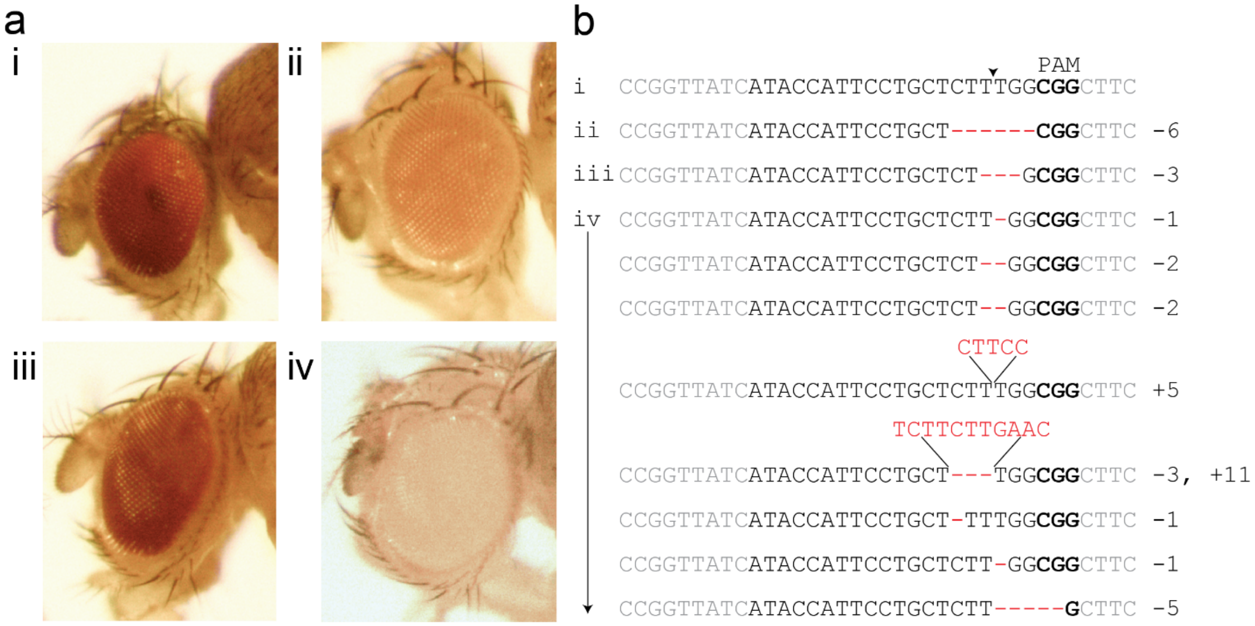


**Supplementary Figure 9: Molecular characterization of CRISPR *white* mutations:** Mutations obtained via CRISPR mutagenesis. (a, I and b, i) Wild-type fly and DNA sequence, (a, ii and b, ii) hypomorphic mutant fly with a 6 bp (non-frameshifting) deletion, (a, iii and b, iii) hypomorphic mutant fly with 3 bp (non-frameshifting) deletion, and (a, iv and b, iv) example image of a null mutant fly and different classes of mutations detected with null mutant phenotypes, including a frameshifting 5 bp insertion, frameshifting 1 bp, 2 bp, and 5 bp deletions, and a complex frameshifting allele carrying a 3 bp deletion coupled with an 11 bp insertion.

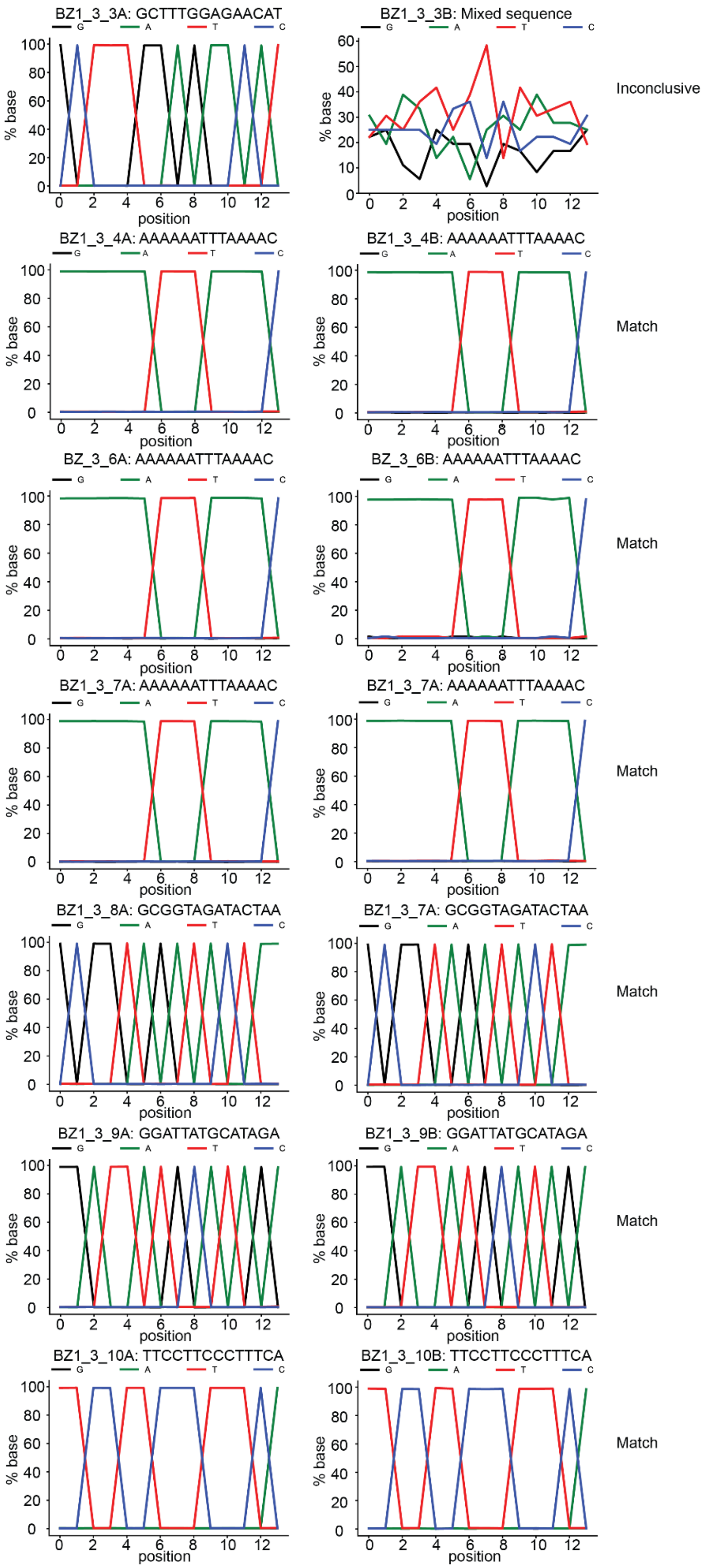
BZ1_3_3A: GCTTTGGAGAACAT
BZ1_3_3B: Mixed sequence
Inconclusive
BZ1_3_4A: AAAAAATTTAAAAC
BZ1_3_4B: AAAAAATTTAAAAC
Match
BZ_3_6A: AAAAAATTTAAAAC
BZ_3_6B: AAAAAATTTAAAAC
Match
BZ1_3_7A: AAAAAATTTAAAAC
BZ1_3_7A: AAAAAATTTAAAAC
Match
BZ1_3_8A: GCGGTAGATACTAA
BZ1_3_7A: GCGGTAGATACTAA
Match
BZ1_3_9A: GGATTATGCATAGA
BZ1_3_9B: GGATTATGCATAGA
Match
BZ1_3_10A: TTCCTTCCCTTTCA
BZ1_3_10B: TTCCTTCCCTTTCA
Match
G
A
T
C
% base
position

**Supplementary Figure 10: Barcode sequencing comparison:** Comparison of replicate barcode lines extracted and sequenced on independent plates.

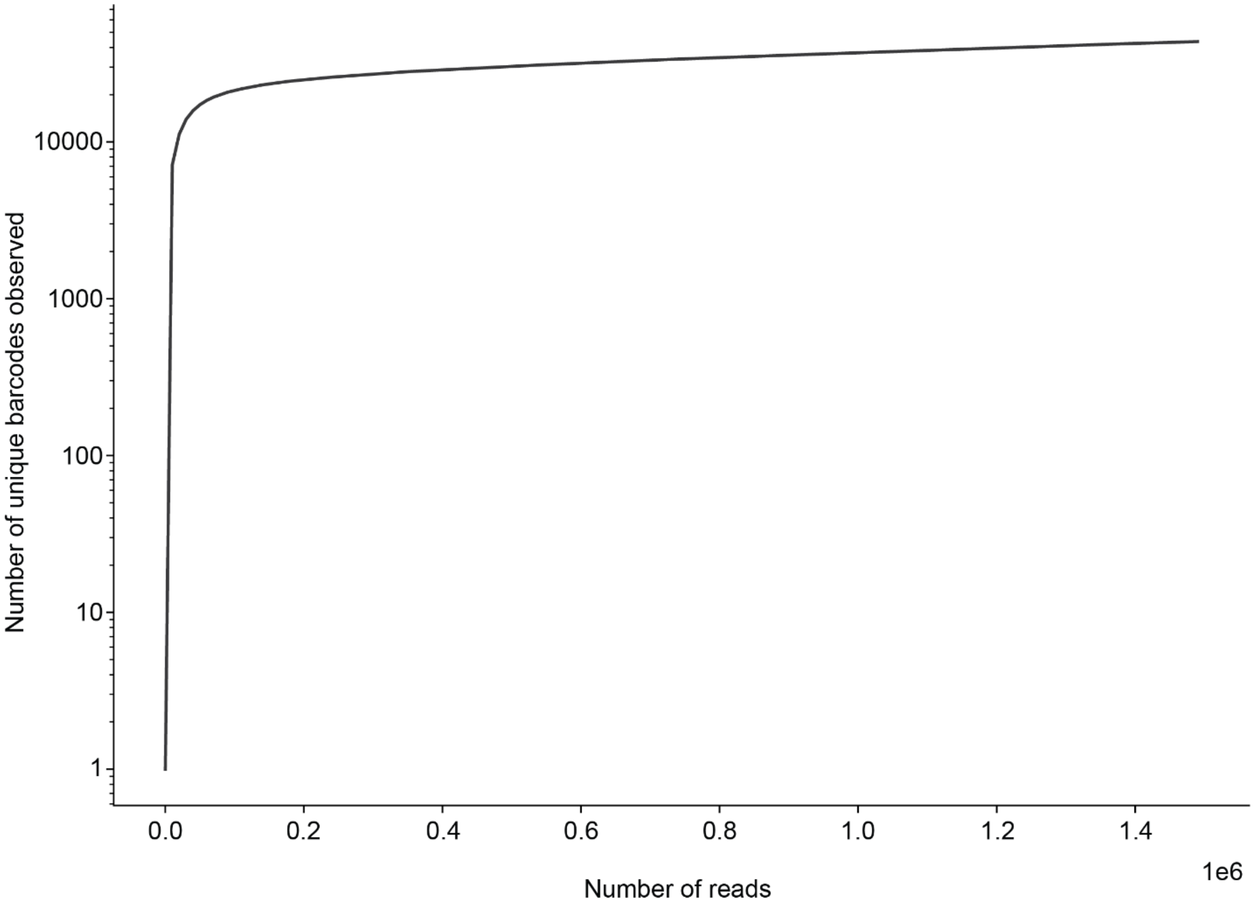


**Supplementary Figure 11: Barcode library saturation:** Barcode library saturation plot.
